# COVID-19 Knowledge, Attitudes, and Practices of United Arab Emirates Medical and Health Sciences Students: A Cross Sectional Study

**DOI:** 10.1101/2021.01.19.427250

**Authors:** Noura Baniyas, Mohamud Sheek-Hussein, Nouf Al Kaabi, Maitha Al Shamsi, Maitha Al Neyadi, Rauda Al Khoori, Suad Ajab, Muhammad Abid, Michael Grivna, Fikri M. Abu Zidan

## Abstract

COVID-19 pandemic is the largest unprecedented viral pandemic of the 21st century. We aimed to study the COVID-19 knowledge, attitudes, and practices (KAP) among medical and health sciences students in the United Arab Emirates (UAE). We performed a cross-sectional study between 2nd June and 19th August 2020. The survey was developed using online Survey Monkey. The link was distributed via UAE University to all students and via WhatsApp© groups. The self-administered questionnaire was conducted in English and comprised of two parts: socio-demographic characteristics and KAP towards COVID-19. A total of 712 responses to the questionnaire were collected. 90% (n=695) were under-graduate, while 10% (n=81) were post-graduate students. Majority (87%, n=647) stated that they obtained COVID-19 information from multiple reliable sources. They were highly knowledgeable about COVID-19 pandemic but 76% (n=539) did not recognize its routes of transmission. 63% (n=431) were worried of getting COVID-19, while 92% (n=633)) were worried that a family member could get infected with the virus. 97% (n=655) took precautions when accepting home deliveries, 94% (n=637) had been washing their hands more frequently, and 95% (n=643) had been wearing face masks. In conclusion, participants showed high levels of knowledge and awareness about COVID-19. They were worried about getting infected themselves or their family members, and had good practices against COVID-19.

## Introduction

COVID-19 is the largest unprecedented viral pandemic of the 21st century. COVID-19 virus (SARS-CoV-2) was first discovered after it caused a cluster of fatal cases of pneumonia in Wuhan, China. The virus has swiftly spread worldwide [1–3]. This pandemic has major impact on global health and economy. Currently there are around 90 million infected persons worldwide, and two million deaths [4–5]. We have to adopt clear effective public health measures so as to mitigate this crisis [6], till we find a proper vaccination [7]. This can not be achieved without evaluating knowledge, attitudes and practices (KAP) towards preventing this disease. This is even more important for role models in the community (like leaders, doctors and medical students) [8]. A study conducted among health care workers in China, during the early stages of the outbreak has shown that around 90% had sufficient knowledge of COVID-19 who followed proper preventative practices. 85% were afraid of getting infected [9]. We thought that it was essential to evaluate the KAP of COVID-19 among medical and health sciences students in the United Arab Emirates (UAE) so to mitigate the effects of this pandemic on them. We aimed to evaluate the knowledge of COVID-19, awareness of preventive behaviors, practice, and risk perception among medical students and allied health sciences in the UAE.

## Materials and methods

### Ethical considerations

Ethical approval was obtained from the UAEU Social Research Committee [UAEU ERS_2020_6119]. Participants’ data were anonymized at the point of registration. No personal identifiable data were collected.

### Study Design

This is a cross-sectional study which was conducted among medical and health sciences students in the UAE between 2^nd^ June and 19^th^ August 2020. The survey was developed online using Survey Monkey. The link was distributed via UAE University to all students and via WhatsApp© groups.

### Sample size

The estimated population of UAE is 9,948,495 [10]. Using Raosoft sample size calculator, having a confidence interval of 95%, a marginal error of 5%, and a response distribution of 50%, the calculated sample size was 385 participants [11]. Accordingly, we aimed to collect 800 participants to assure reaching our objectives.

The online questionnaire was aimed to reach the target population. We used the snowball sampling technique. The study invitation and survey link were directly sent to the medical and health sciences colleges and universities in the UAE through the e-mail and circulated on multiple social media outlets. The participants were encouraged to forward the link to their fellow medical and health sciences students and post it on their social media platforms to maximize enrolment of potential participants. The study invitation included an introduction, a brief description of the study and the link to the questionnaire. The survey was piloted on 10 students and validated by epidemiologists before sharing it with the study population.

### Questionnaire design

The questionnaire was designed and referenced from previous similar studies and modified to our study population [12,13]. The questionnaire started with a consent page which provided a brief description of the study, the voluntary nature of participation and declaration of anonymity. Online informed consent was obtained prior proceeding to the questionnaire response. The self-administered questionnaire was conducted in English and comprised of two parts: socio-demographics characteristics and KAP towards COVID-19. The demographic variables included age, gender, nationality, place of residence, if they are currently studying medicine or health sciences, their year of study and specialty, if they suffer from chronic diseases or live with someone who has a chronic disease, if they tested positive for COVID-19, their source of COVID-19 information, and if they attended any COVID-19 educational courses. The KAP part consisted of 3 sections and a total of 34 questions. These included

#### Knowledge

This section included 12 questions which assessed the participants’ knowledge about COVID-19, and the questions were answered on a multiple choice and true/false basis. The items included etiology of the disease, transmission of the virus, symptoms, incubation period, diagnostic tests, treatment options, and prevention. In the knowledge section, respondents were given options to answer as true, false, or don’t know.

#### Attitude

This section included 6 questions which assessed the participants’ attitudes towards the COVID-19 pandemic using a Likert scale. This was coded as follows: strongly disagree=1, disagree=2, undecided=3, agree=4, strongly agree=5. This included fear of getting infected, stigma around infected individuals, government measures, and their confidence towards the measures.

#### Practice

This section included 16 questions which assessed the participants’ practices towards COVID-19 using multiple-choice questions, yes/no questions, and a Likert scale. The items were related to practices and compliance to preventative measures and precautions implemented by the government such as social distancing, wearing face masks and hand washing.

### Statistical Analysis

Simple summary descriptive statistics were used. Continuous were presented as mean (SD) while categorical data were presented as number (%). Percentages were calculated from actual available responses. We used the Statistical Package for the Social Sciences (IBM-SPSS version 26, Chicago, Il) for statistical analysis.

## Results

A total of 712 responses to the questionnaire were collected. **Table 1** shows the detailed demography of the participants. 90% (n=695) of respondents were under-graduates, while 10 % (n=81) were post-graduate. The majority of respondents (87%, n=647) stated that they obtained COVID-19 information from multiple sources. Social Media was the main source for COVID-19 information (7%, n=52) of the respondents while the rest (6%, n=48) relied on medical platforms, healthcare professionals, government media briefings, and university newsletters. 406 respondents (57%) attended webinars to learn more about COVID-19.

**Table 1.**
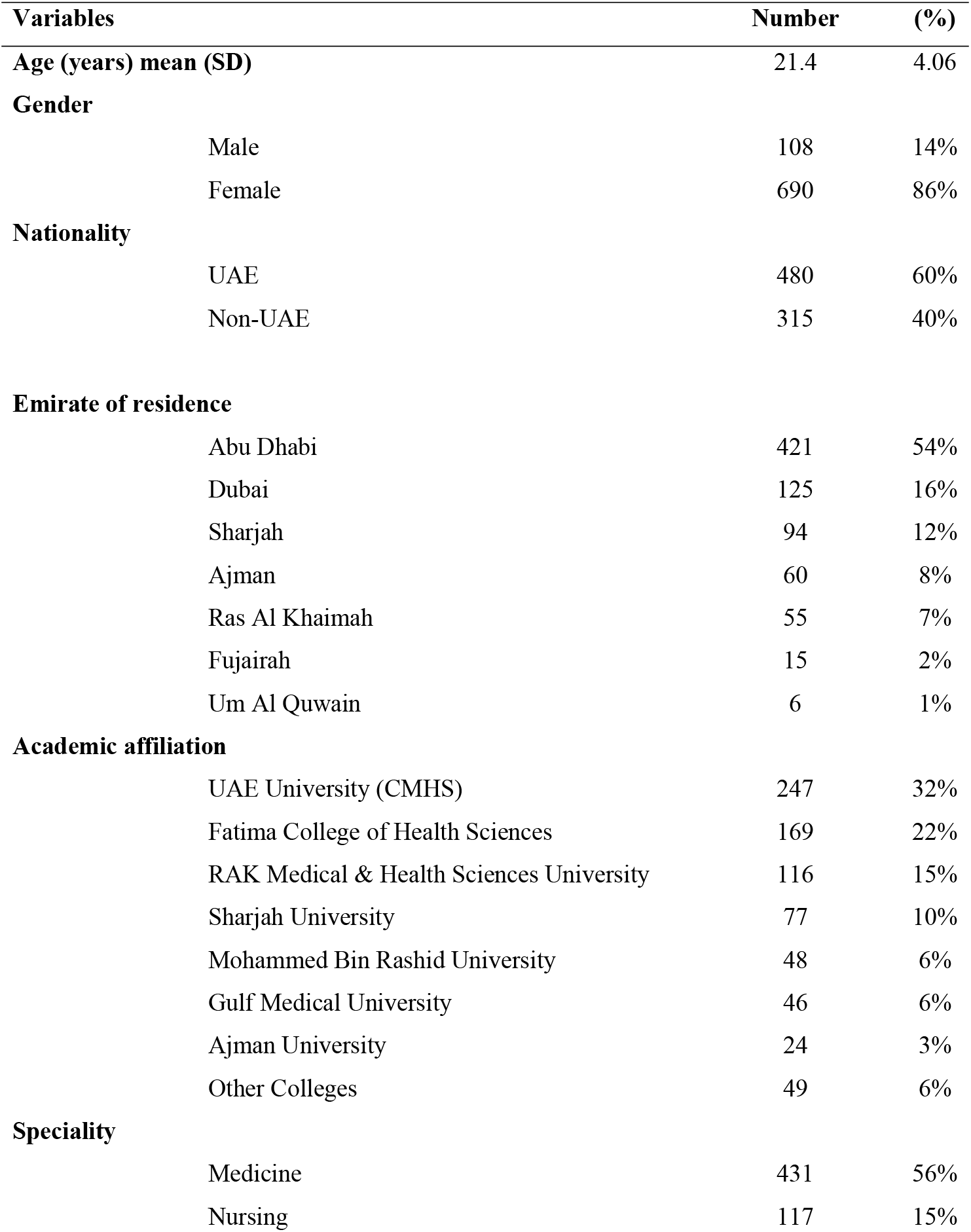
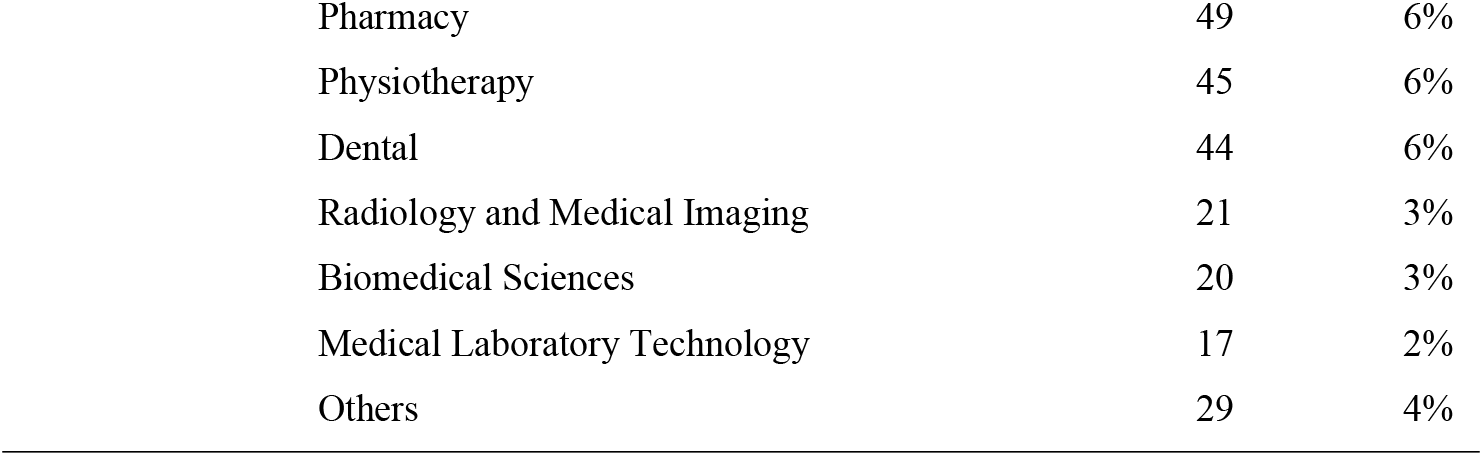
Characteristics of respondents of the KAP survey collected between 2^nd^ June and 19^th^ August 2020

| **Variables** |  | **Number** | **(%)** |
| --- | --- | --- | --- |
| **Age (years) mean (SD)** |  | 21.4 | 4.06 |
| **Gender** |  |  |  |
|  | Male | 108 | 14% |
|  | Female | 690 | 86% |
| **Nationality** |  |  |  |
|  | UAE | 480 | 60% |
|  | Non-UAE | 315 | 40% |
| **Emirate of residence** |  |  |  |
|  | Abu Dhabi | 421 | 54% |
|  | Dubai | 125 | 16% |
|  | Sharjah | 94 | 12% |
|  | Ajman | 60 | 8% |
|  | Ras Al Khaimah | 55 | 7% |
|  | Fujairah | 15 | 2% |
|  | Um Al Quwain | 6 | 1% |
| **Academic affiliation** |  |  |  |
|  | UAE University (CMHS) | 247 | 32% |
|  | Fatima College of Health Sciences | 169 | 22% |
|  | RAK Medical & Health Sciences University | 116 | 15% |
|  | Sharjah University | 77 | 10% |
|  | Mohammed Bin Rashid University | 48 | 6% |
|  | Gulf Medical University | 46 | 6% |
|  | Ajman University | 24 | 3% |
|  | Other Colleges | 49 | 6% |
| **Speciality** |  |  |  |
|  | Medicine | 431 | 56% |
|  | Nursing | 117 | 15% |

|  |  |  |
| --- | --- | --- |
| Pharmacy | 49 | 6% |
| Physiotherapy | 45 | 6% |
| Dental | 44 | 6% |
| Radiology and Medical Imaging | 21 | 3% |
| Biomedical Sciences | 20 | 3% |
| Medical Laboratory Technology | 17 | 2% |
| Others | 29 | 4% |

**Table 2** shows the comorbidities and COVID-19 history of the participants. 8% of the participants who were tested for COVID-19 were positive. 85% (n=506) had a family member or friend who got tested for COVID-19, of which 15% (n=89) tested positive.

**Table 2.** Comorbidities and COVID-19 history of the KAP survey respondents

| Variable | n | (%) |
| --- | --- | --- |
| **Personal chronic condition** |  |  |
| Asthma | 28 | 37% |
| Diabetes | 10 | 13% |
| Hypertension | 5 | 7% |
| Inflammatory bowel disease | 3 | 4% |
| Migraine | 3 | 4% |
| Polycystic ovary syndrome | 3 | 4% |
| Others | 24 | 32% |
| **Personal history of COVID-19** |  |  |
| Tested for COVID-19= yes | 160 | 92% |
| Tested positive= yes | 13 | 8% |
| **Household history of COVID-19** |  |  |
| Asymptomatic | 16 | 19% |
| Quarantined with mild symptom | 57 | 67% |
| Admitted to hospital with severe symptoms | 6 | 7% |
| Admitted to ICU with severe symptoms | 3 | 4% |
| Died | 3 | 4% |

### Knowledge

A total of 712 respondents completed the knowledge section of the survey (Table 3). 76% (n=539) of participants did not recognize the correct routes of transmission of COVID-19, while the majority of respondents correctly recognized its symptoms, average incubation period, best diagnostic test, and its management (95%, 85%, 89%, 89% and 70% respectively). The majority of the respondents were aware of the COVID-19 preventative measures including methods to reduce viral spread, isolation of positive cases, N95 mask use limited to health care workers, and the necessity of preventative precautions among young adults and children (83%, 92%, 84%, and 87% respectively).

**Table 3.** Responses to the survey on COVID-19 knowledge

| Statement | Correct |  | Incorrect/<br>Uncertain |  | Total* |
| --- | --- | --- | --- | --- | --- |
|  | n | (%)* | n | (%)* |  |
| COVID-19 is a new disease caused by virus SARS-CoV-2. | 570 | 80% | 142 | 20% | 712 |
| Which animal is most likely to transmit this virus to human? | 633 | 89% | 79 | 11% | 712 |
| SARS-CoV-2 can be transmitted between humans by the following routes? | 173 | 24% | 539 | 76% | 712 |
| Which of the following are COVID-19 symptoms? | 679 | 95% | 33 | 5% | 712 |
| What is the average incubation period of COVID-19? | 602 | 85% | 110 | 15% | 712 |
| What is the best diagnostic test for COVID-19? | 636 | 89% | 76 | 11% | 712 |
| COVID-19 can be treated by using the following: | 500 | 70% | 212 | 30% | 712 |
| Which of the following can reduce the spread of COVID-19? | 591 | 83% | 121 | 17% | 712 |
| People who are asymptomatic and COVID-19 test positive must stay at home until they are free of the infection: | 653 | 92% | 59 | 8% | 712 |
| Who should wear N95 masks? | 601 | 84% | 111 | 16% | 712 |
| Persons with COVID-19 cannot transmit the virus to others when a fever is not present. | 635 | 89% | 77 | 11% | 712 |
| It is not necessary for children and young adults to take measures to prevent infection from COVID-19 virus. | 616 | 87% | 96 | 13% | 712 |

Percentages may not total 100 because of rounding
\*Percentages were calculated out of 712, which is the total number of respondents who completed the knowledge items

### Attitude

A total of 686 respondents completed the attitudes section of the survey (**Table 4**). 63% (n=431) of participants were worried of getting COVID-19 infection, while the vast majority (92% (n=633)) were worried that a family member could get infected with the virus. 67% (n=461) of the respondents thought that infection with the virus is associated with stigma. 83% (n=570) agreed that the current measures taken by the UAE government are effective in stopping the spread of the infection and 89% (n=614) were confident that the UAE will be able to stop the spread of the virus. Nevertheless, 60% (n=288) thought that more measures could be implemented such as aggressive screening, full lockdown, further education to the public, monitoring the media, and fighting rumors. Some went against the lockdown and suggested to loosen the restrictions.

**Table 4.**
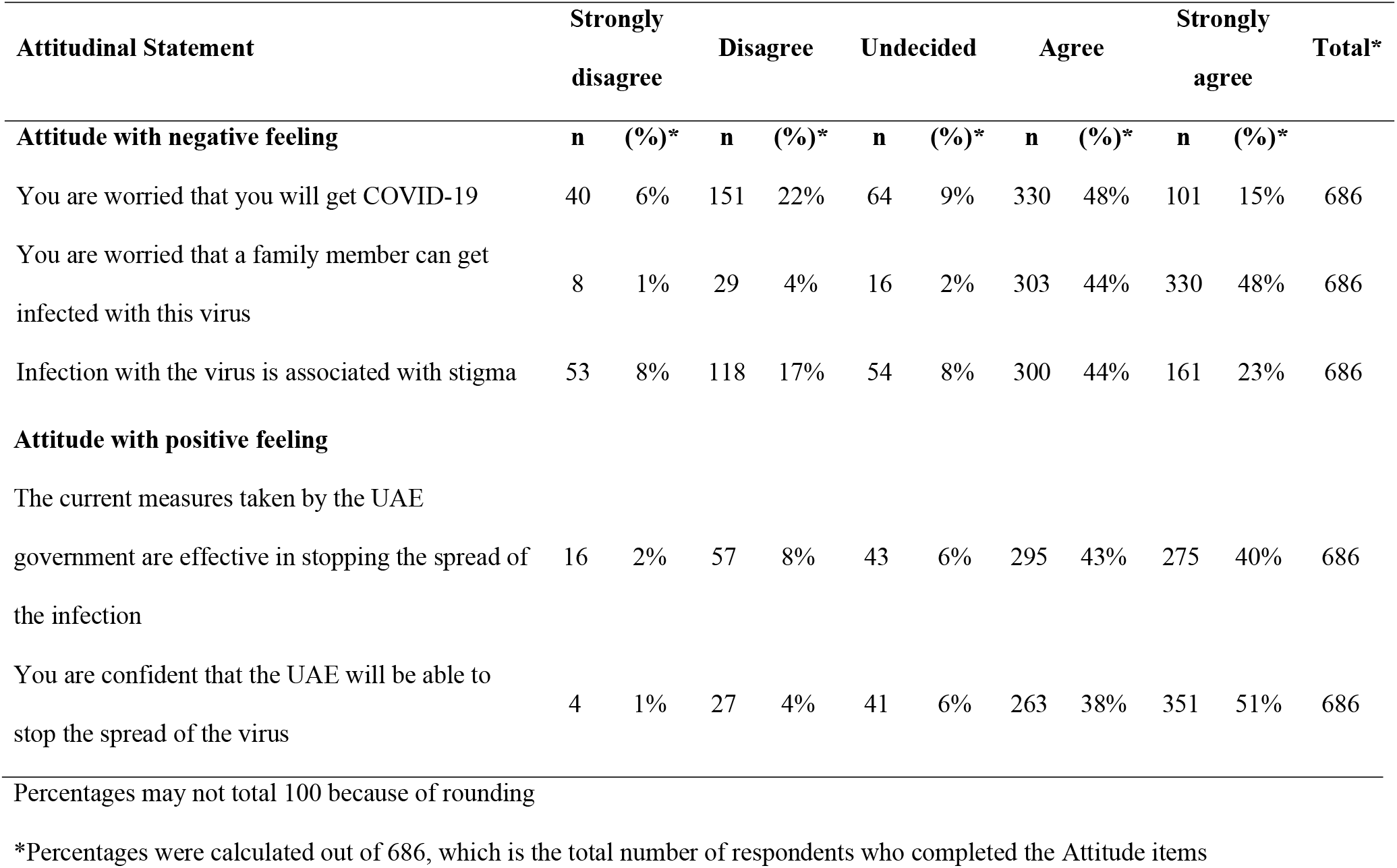
Responses to the survey on COVID-19 Attitude

### Practices

A total of 677 respondents completed the practices section of the survey. 60% (n=407) did not attend family gatherings, did not visit shopping malls, coffee shops, industrial areas, hospitals or COVID-19 facilities for volunteering. 97% (n=655) took precautions when accepting home deliveries, 94% (n=637) had been washing their hands more frequently, and 95% (n=643) had been wearing face masks. Concurrently, out of 666 respondents, nearly all of them followed curfew timings set by the UAE government (99% (n=658)). Overall, most medical students and allied health sciences students followed proper practices.

**Table 5.**
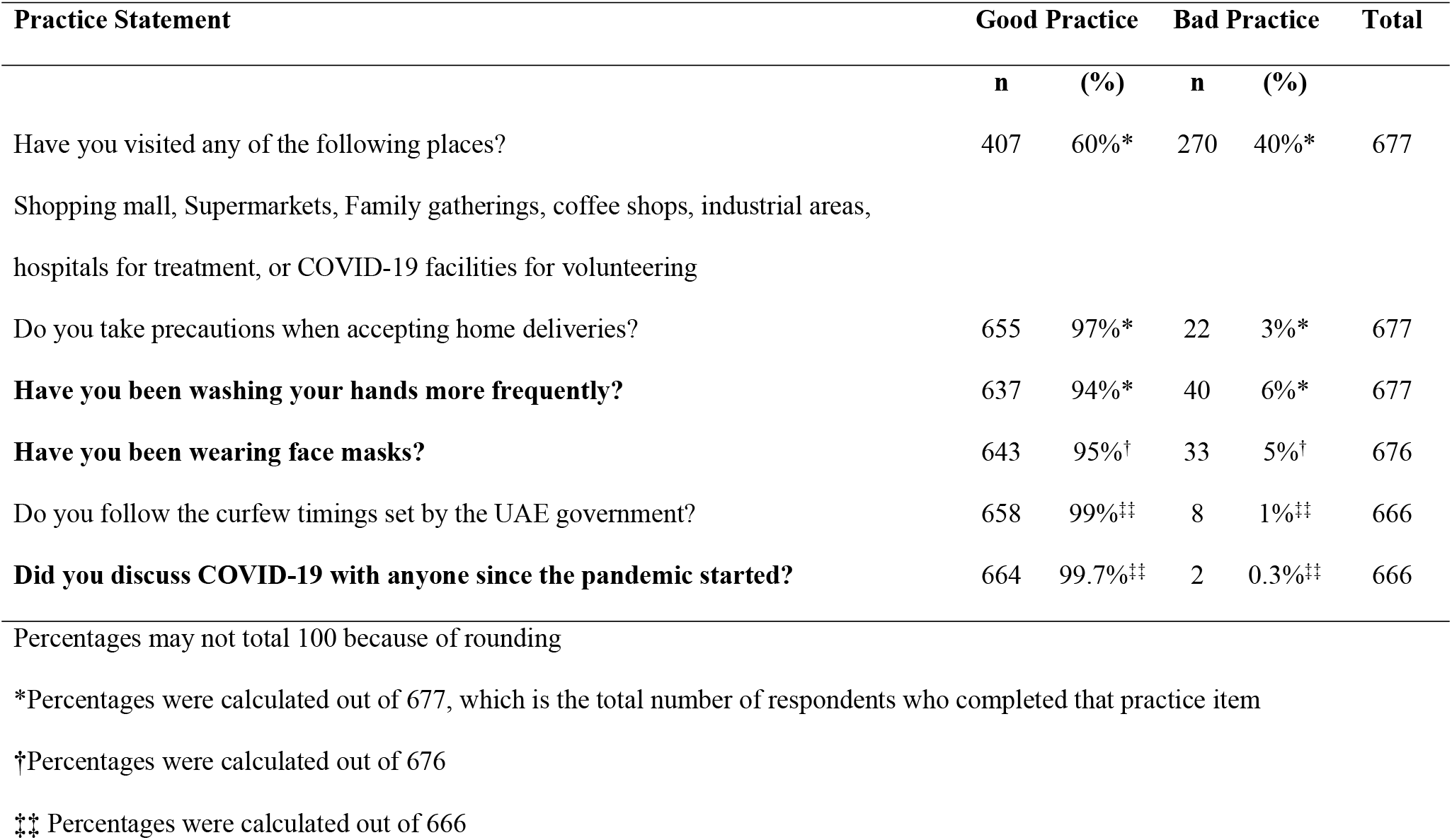
Responses to the survey on COVID-19 Practices

## Discussion

Our study has shown that the majority of medical and allied health students at UAE were well knowledgeable about COVID 19, worried about getting infected or having a member of their family infected, and had proper practices and precautionary measures for preventing COVID-19.

The high knowledge on COVID-19 of the medical and allied health students in the UAE is similar to those reported from Pakistan [14]. This may be attributed to the use of multiple reliable medical platforms, healthcare professionals, government media briefings, and university newsletters. These resources may have enforced proper knowledge. Furthermore, it may be related to the training they received as volunteers in the healthcare system [15]. The majority of the respondents were aware of COVID 19 symptoms, the incubation period, diagnostic testing, management, and the preventative measures. However, only 24% in our study correctly recognized COVID-19 route of transmission compared with other studies in which undergraduate students were quite knowledgeable about the route of transmission [14, 16]. COVID-19 is transmitted by respiratory droplets; however, airborne transmission may be possible in case of a medical procedure that generates aerosols [17]. This was unexpected, given the fact that majority of our respondents depended on reliable medical resources for their information.

Furthermore, our study has shown that participants had both negative and positive feelings regarding the pandemic. Majority was worried that they or a member of their family may get infected. It is important to note that this study was conducted during the period of the rapid increase of COVID-19 cases in the UAE. 67% of the participants believed that infection with the virus is associated with stigma. This may lead to high-risk behaviors that increase infection such as gatherings, poor hand hygiene, poor social distancing and not following the government guidelines. Interestingly, most of the participants were confident that the UAE will be able to stop the spread of the virus. They also believed that the current measures taken by the UAE government are effective in stopping the spread of the infection. This positive attitude can be explained by the drastic measures taken by the UAE government to contain the spread of the virus. These measures included the introduction of teleworking, distance learning, lockdowns, country-level coordination, risk communication, community engagement, surveillance, rapid response teams, case investigation, infection prevention and control, and operational support and logistics [18].

Majority of participants had proper practices and precautionary measures against COVID-19. 60% did not visit shopping malls, family gatherings, coffee shops, industrial areas and hospitals; and the majority reported good preventative practices including hand washing, wearing face masks, and abiding by curfew timings. These findings could be attributed to the strict lockdown when the survey was launched, access to trusted medical resources, and training in medical fields. These results are similar to those reported on undergraduate students in China, medical students in Pakistan, and clinical year medical students in Iran [13–16].

### Limitations

We have to acknowledge that out study has certain limitations. First, this study was conducted using an online survey; consequently, the results of the questionnaire all depended on the participants self-reported behaviors, with no means of confirming whether the responses were exaggerated as a result of social desirability bias. Second, the respondents were predominantly females and medical students. This may be a selection bias with its effect on the results. Finally, this study was conducted in the early stages of the pandemic when the UAE was under lockdown and continued for a while after the restrictions were lifted. Since then, more information about the pandemic has been published and likewise public health measures in the UAE have changed. Thus, the results of the study may not represent the current COVID-19 KAP of the medical and health sciences students.

## Conclusions

Medical and health sciences students in the UAE showed high levels of knowledge and awareness about COVID-19. Although they were confident in the public health measures taken to mitigate the COVID-19 pandemic, they were worried about getting infected themselves or their family members, and had good practices against COVID-19.

## Abbreviations

SARS-CoV-2: Severe acute respiratory syndrome coronavirus 2
COVID-19: Coronavirus disease 2019
HCW: Healthcare workers
ICU: Intensive care unit
ILO: International Labour Organization
IFAD: The International Fund for Agricultural Development
WHO: World health organization
FAO: Food and Agriculture Organization
KAP: knowledge, attitudes, and practices
UAE: United Arab Emirates
UAEU: United Arab Emirates University

## Declarations

### Ethics approval

Ethical approval was obtained from the UAEU Social Research Committee (UAEU ERS_2020_6119).

### Consent for publication

Not applicable

### Availabilities of data and material

Data will be available as excel file after acceptance of the paper.

### Contribution of authors

Noura Baniyas, Rauda Al Khoori, Maitha Al Shamsi, Maitha Al Neyadi, Nouf Al Kaabi, and Suad Ajab formulated the research question, developed the protocol, collected and coded the data, and wrote the first draft. Michal Grivna and Muhammad Abid contributed to the questionnaire design and revised the first manuscript. M. Sheek-Hussein, Supervised, the project including the questionnaire design, the analysis and drafting of the first version of the paper. Fikri Abu Zidan, supervised the analysis of the data of the questionnaire and writing the first version of the paper, and extensively edited the paper. All authors contributed, revised the final manuscript, and approved.

## Acknowledgements

We are thankful to Dr. Ahmed R. Alsuwaidi, Dr. Iffat Elbarazi, Laila Masood, Ph.D. Candidate, Dr. Marilia Silva Paulo for their advice during developing this project, and for Ms. Laila Masood for facilitating the SurveyMonkey.

